# A constraints-based theory of the primary cause of senescence: imbalance of epigenetic and non-epigenetic information in histone crosstalk

**DOI:** 10.1101/310300

**Authors:** Felipe A. Veloso

## Abstract

Cellular aging has been progressively elucidated by science. However, the fundamental cause of senescence—i.e., why organisms age at the multicellular-individual level—remains unclear. A recent theory of individuated multicellularity describes the emergence and growth of crucial information content for cell differentiation. This information is mostly conveyed in the non-epigenetic (i.e., transcription uncorrelated) histone crosstalk near transcription start sites. According to this theory, the non-epigenetic content emerges and grows at the expense of the information capacity for epigenetic content. If this “reassignment” of information capacity continues after adulthood, it may explain the senescence phenomenon. Here, I present a novel, falsifiable theory describing an uninterrupted growth of capacity for non-epigenetic information at the expense of that for epigenetic information not only during ontogeny but also throughout adulthood. As a byproduct, this continuous “reassignment” of capacity effectively creates an information imbalance in histone crosstalk, which in turn overregulates transcriptional levels. This overregulation is to be understood as transcriptional levels becoming more and more accurate but also less and less precise with respect to the needs of the multicellular individual—up to the point of dysfunctionality. This epigenetic/non-epigenetic information imbalance is proposed to be the primary reason why individuated multicellular organisms senesce.

## 1. INTRODUCTION

### 1.1. Background

Our intellectual endeavors have entertained the prospect of unlimited lifespan for centuries^1^, and the scientific endeavor has been no exception^2^. In the 1950s, the immortality of cultured somatic cells was indeed a widely-held belief^3^. That changed only when Hayflick & Moorhead showed that cultured human somatic cells do stop dividing and become less viable once their divisions reach a certain number^4^, a phenomenon known today as the Hayflick limit^3^. This loss of replicative capacity and, in general, the process of aging at the cellular level, have been elucidated to a very considerable extent^5^ (and references therein). Yet, the number of times human cells can divide in culture exceeds the number of times cells divide throughout our lifespan; there is no significant correlation between human cell replicative capacity and cell donor age^6^. That is, we—and individuated multicellular organisms in general—senesce (see Glossary) before most of our cells do^7,8^.

### 1.2. Scope

The theory presented here is not aimed to explain the complex cascade of dysfunctional changes that characterizes the senescence of a multicellular individual^8^ (and references therein) but to provide a falsifiable explanatory account of the very beginning of said cascade in terms of Mayr’s proximate cause and ultimate cause^9^. In other words, this theory pertains to *how* the senescence process is triggered throughout the adulthood of a multicellular individual (i.e., proximate cause) and *why*, in evolutionary terms, most multicellular species senesce (i.e., ultimate cause).

### 1.3. Senescence’s accepted ultimate cause: a critique

Senescence is widely regarded as an evolutionary consequence of the relaxation of selection on traits that maintain/repair the multicellular individual’s functions in later life, because later life would have been rarely realized in the wild with the hazards it imposes^10–12^. Yet, any well-established ultimate cause of senescence must seamlessly integrate with the associated proximate cause, which itself remains unclear.

Notwithstanding, efforts have been made to develop an explanatory account that integrates relaxed selection as its ultimate cause with a suggested proximate cause.

The first is the explanatory account dubbed as “disposable soma”. This account states that somatic maintenance requires an amount of resources that are often very limited and thus require trade-offs between the components of maintenance (growth, reproduction, and self-repair) in order to maximize evolutionary fitness^13,14^. Here the proximal cause is not fully described but it entails the accumulation of damage over time (somehow leading to senescence) when insufficient resources are spent on self-repair. According to this account, an individual will spend the minimal amount of resources sufficient to reach sexual maturity, after which its soma becomes disposable^13,14^.

A number of objections have been raised in this regard, such as the fact that variations in reproduction and lifespan are not always associated—as they should be, according to the disposable soma account^15^. Among mammals, naked mole-rats exemplify that reproductive and non-reproductive individuals can display similar maximum lifespans, i.e., no observable association^16^.

Recapitulating, it is safe to observe that (i) the proximate cause of senescence is fundamentally unclear (e.g., we remain ignorant of how species such as ours begin to senesce while a few others such as the naked mole-rat display negligible senescence^17^) and, for this very reason, (ii) relaxed selection being the ultimate cause of senescence should not be, at least up to now, accepted as a well-established scientific fact.

### 1.4. Alternative proximate causes

Other proposed proximate causes of senescence or aging at the multicellular-individual level have been classified into two categories: programmed senescence and senescence caused by damage/error^18^. Recently it has been argued, however, that senescence is not programmed nor is it ultimately a consequence of damage or error in the organism’s structure/dynamics^19^. Instead, it may be a byproduct of maintenance and/or developmental dynamics^19,20^, themselves underpinned in part by intracellular signaling pathways such as the cell-cycle-related PI3K/AKT/mTOR pathway^19^. Some of these pathways have been shown to modulate aging at the cellular level in species such as the yeast *Saccharomyces cerevisiae*^21^.

The analogous notion of aging at the multicellular-individual level as a byproduct of certain functional signaling pathways^19^ is, in principle, supported by the fact that a mutation in the *daf-2* gene—which encodes the insuline-like growth factor 1 receptor in the PI 3-kinase pathway—and the deficiency of mTOR kinase—a key component of the PI3K/AKT/mTOR pathway—can double the lifespan of the roundworm *Caenorhabditis elegans*^22,23^. The hypotheses where senescence is the effect of a runaway activity of processes useful in young age that have become harmful in old age have been dubbed “bloated soma” hypotheses^24^.

Alas, the fundamental dynamics that trigger senescence in individuated multicellular organisms remain unclear and most explanatory accounts^18,25^ (and references therein) lack predictions to falsify them.

### 1.5. Towards a constraints-based theory of senescence

Here, I suggest that senescence is a dysfunctional byproduct of functional developmental dynamics as first described by a recently proposed theory of individuated multicellularity^26^. Specifically, I show that the byproduct is a post-ontogenetic, growing imbalance between two different information contents conveyed respectively in two different types of constraints on histone post-translational modifications nearby transcription start sites (TSSs). Constraints are here understood as the local and level-of-scale specific thermodynamic boundary conditions required for energy to be released as work as described by Atkins^27^. The concept of constraint is crucial because, according to the theory of individuated multicellularity, a higher-order constraint (i.e., a constraint on constraints) on changes in histone modifications harnesses critical work that regulates transcriptional changes for cell differentiation at the multicellular-individual level.

Under the theory of individuated multicellularity, the intrinsic higher-order constraint is the simplest multicellular individual in fundamental terms. In addition, the dynamics of the lower-order constraints must be explicitly unrelated to each other in order to elicit the emergence of the intrinsic higher-order constraint^26^ (i.e., statistically independent). Along with the emergence of this intrinsic higher-order constraint, the theory of individuated multicellularity describes the emergence of critical information content, named in the theory hologenic content, which is about the multicellular individual as a whole in terms of developmental self-regulation. Thus, for the sake of brevity, I here refer to the theory of individuated multicellularity as the hologenic theory.

The constraints on the combinatorial patterns of nucleosomal histone post-translational modifications (PTMs) are generally known as histone crosstalk^28,29^. Histone modifications are also known to be relevant for epigenetic changes^30^, which are defined as changes in gene expression that cannot be explained by (i.e., that are explicitly unrelated to) changes in the DNA sequence^31^. Moreover, specific histone modifications are known to be associated with senescence^32^ (and references therein). These associations are underpinned by the ability of histone modifications to convey information content, which has allowed the prediction of mRNA levels from histone modification profiles nearby TSSs with high accuracy^33^.

**Glossary**
senescence / biological aging: time-dependent, progressive impairment of biological functions undergone by individuals of most multicellular species once they reach their mature form
cellular senescence / cellular aging: irreversible arrest of cell proliferation; this phenomenon can be observed in the laboratory once somatic, differentiated cells reach a number of divisions known as Hayflick limit
accuracy: average closeness of a number of measured values to a target value
precision: variance of a number of measured values
work: constrained release of energy (see ref. 27)
constraints: local and level-of-scale specific thermodynamic boundary conditions
histone crosstalk: combinatorial patterns of histone post-translational modifications
information content: emergent, generative constraints on the dynamics of a living system resulting from energy released by it under constraints already embodied in a physical medium (see refs. 68 and 69)
information capacity: amount of information a physical medium (e.g., histone crosstalk) *can* embody, typically expressed in bits
total correlation: multivariate generalization of Shannon’s mutual information, useful for measuring information capacity in histone crosstalk beacause of its mathematical properties (see refs. 34 and 35)

Based on these considerations and the properties of the non-negative measure of multivariate statistical association known as total correlation^34^ or multiinformation^35^ (symbolized by *C* and typically measured in bits), the overall observable histone crosstalk can be decomposed. That is, histone crosstalk, if measured as a total correlation *C*, is the sum of two explicitly unrelated *C* components: one epigenetic (i.e., explicitly related to changes in gene expression) and the other non-epigenetic (i.e., explicitly unrelated to changes in gene expression). This sum can be expressed as follows:

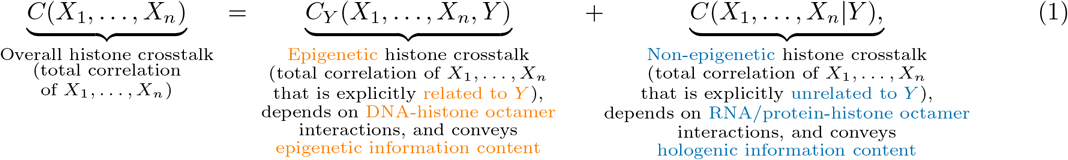

where *X*_1_,…, *X_n_* are random variables representing *n* histone modification levels in specific genomic positions with respect to the TSS and *Y* is a random variable representing either gene expression level, transcription rate, or mRNA abundance level associated with the TSS. These levels are equivalent for the decomposition because of the strong correlation that exists between them^36^ (and references therein).

Importantly, the three terms in Eq. 1 correspond to statistical associations between the random variables *X*_1_,…, *X_n_*, *Y* (histone PTM and mRNA levels) that take their respective values from data for all transcriptional start sites. Therefore, these terms must be understood as representing chromatin-wide constraints.

The hologenic theory describes how the epigenetic component of histone crosstalk (represented by *C_Y_* (*X*_1_,…, *X_n_, Y*) in the sum decomposition of Eq. 1) conveys information about each cell’s transcriptional profile. This component is, in information content terms, the dominating component for any eukaryotic colonial species—such as the alga *Volvox carteri*^37^—and, importantly, also for undifferentiated stem cells.

The second, non-epigenetic component of histone crosstalk (represented by *C*(*X*_1_,…, *X_n_*|*Y*) in Eq. 1) is known to grow in magnitude during development until the organism’s mature form is reached^26^. This component is described by the hologenic theory as conveying information about the multicellular individual as a whole—starting from the moment said individual emerges as an intrinsic higher-order constraint on the early embryo’s proliferating cells.

Importantly, the overall observable histone crosstalk magnitude (represented by *C*(*X*_1_,…, *X_n_*) in Eq. 1) is not infinite. In other words, the overall histone crosstalk has a finite information capacity, which can be measured in bits. Moreover, the sum decomposition in Eq. 1 implies that, for a given (constant) overall histone crosstalk, the growth in magnitude (bits) of the hologenic (i.e., non-epigenetic) component must be accompanied by a decrease in magnitude of the epigenetic component. That is, the capacity (in bits) for hologenic information content in histone crosstalk is bound to grow at the expense of the capacity for epigenetic information content.

The hologenic theory also maintains that a necessary condition for the evolution of individuated multicellular lineages was the appearance of a class of molecules synthesized by the cells—called Nanney’s extracellular propagators (symbolized by 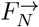) in the theory^26^. These 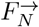 molecules are predicted to be, in a given tissue and time period, (i) secretable into the extracellular space, (ii) once secreted, capable of eliciting a significant incremental change (via signal transduction) in the magnitude of the non-epigenetic histone crosstalk (i.e., the *C*(*X*_1_,…, *X_n_*|*Y*) summand in Eq. 1) within other cells’ nuclei, and (iii) affected in their extracellular diffusion dynamics by the geometrical complexity of the extracellular space (i.e., constraints on diffusion at the multicellular-individual level, which cannot be reduced to constraints at the cellular level). Also under the hologenic theory, for the multicellular individual to develop and survive, both hologenic (developmental self-regulation of the multicellular individual overall) and epigenetic (each cell’s transcriptional profile) contents must coexist.

One final but important consideration regarding histone crosstalk is that it is the result of constraints which, as mentioned previously, are level-of-scale specific. To exemplify this specificity, consider an internal combustion engine: a single molecule in a cylinder wall or the entire engine do not embody a constraint on the expansion of the igniting gas, yet the cylinder-piston ensemble does. For this reason, histone crosstalk constraints are expected to have relevance for senescence but only at a specific level of scale.

To investigate from a theoretical standpoint if the “reassignment” of information capacity for epigenetic and non-epigenetic (i.e., hologenic) content stops when development reaches the multicellular individual’s mature form or instead continues without interruption, one also needs to investigate the “reassignment” (if any) in cancer cells. One of the corollaries of the hologenic theory is a significant loss of hologenic content in cancer cells, because they are no longer constrained by the multicellular individual that normal (i.e., non-cancerous) cells serve and are constrained by. Thus, I developed a falsifiable theory of senescence based on the post-ontogenetic continuation of this “reassignment” process in histone crosstalk.

### 1.6. How to measure histone crosstalk

Publicly available high-throughput data of histone H3 modifications—because of their high predictive power on transcriptional levels^33^—and mRNA abundance in primary cells can be used to formalize a theoretical description to the senescence phenomenon.

Using these tandem ChIP-seq and RNA-seq data, one can quantify the epigenetic and non-epigenetic (i.e., hologenic) capacities in the crosstalk within a core histone such as the H3 histone (Eq. 1) for some specific level of scale, e.g., for triads of variables {*X_i_, X_j_, X_k_*}. These variables represented position-specific histone H3 modification levels, i.e., *C*(*X_i_, X_j_, X_k_*|*Y*) and *C_Y_*(*X_i_, X_j_, X_k_, Y*) for the non-epigenetic and epigenetic histone crosstalk components, respectively, where *Y* represents mRNA abundance.

The log-ratio between the non-epigenetic and epigenetic histone H3 crosstalk magnitudes can be thus computed as the dimensionless quantity

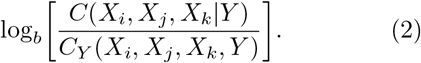

Importantly, this log-ratio derives from the right-side terms of Eq. 1, therefore it must be understood as a chromatin-wide relationship. (Note: total correlation *C* captures all possible associations in the set of (in this example, three) variables {*X_i_, *X_j_*, X_k_*} that may exist starting from the pairwise level. When *b*=2 the information capacity represented by total correlation *C* is measured in bits.)

## 2. A CONSTRAINTS-BASED THEORY OF SENESCENCE

### 2.1. Characterizing the information capacity “reassignment” in histone crosstalk

At some level of scale—without loss of generality we will assume such level comprises up to triads of variables—for normal primary somatic cells of the same type there is a positive and highly significant correlation between the hologenic/epigenetic log-ratio quantity in Eq. 2 and the age of the multicellular individual:

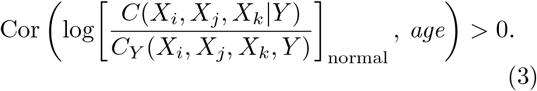

On the other hand, at the same level of scale the positive correlation in Eq. 3 exists, for cancer cells the correlation between the hologenic/epigenetic log-ratio in Eq. 2 and the age of the multicellular individual is not statistically significant:

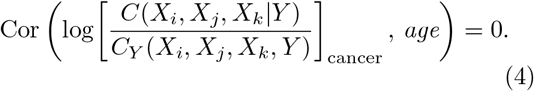

### 2.2. Senescence as transcriptional overregulation

This age-correlated hologenic/epigenetic information imbalance in histone crosstalk can also be understood in terms of an imbalance between the accuracy and precision of transcription in the cells with respect to the needs of the multicellular individual.

If transcriptional regulation of any given gene in any given is understood as a constraint on its associated mRNA levels such that they are on average close to a certain target value (i.e., making said mRNA levels accurate), then under this theory transcription becomes “overregulated” with age in the sense that more accuracy is gained at the increasing expense of precision (i.e., closeness of the resulting mRNA levels to their own mean). This accuracy-precision trade-off—see sum decomposition in Eq. 1—is unavoidable because (i) the relative growth of *C*(*X*_1_,…, *X_n_*|*Y*) implies an increasing constraint on (i.e., regulation of) histone modification patterns with respect to the multicellular individual^26^, thus making transcription more accurate and (ii) the concurrent relative decrease of *C_Y_*(*X*_1_,…, *X_n_, Y*) means histone modification patterns become worse and worse predictors of mRNA levels, in turn making transcription less and less precise up to the point of dysfunctionality with respect to the multicellular individual (see schematic in Fig. 1a). Importantly, this theoretical description parsimoniously explains the age-dependent decrease in transcriptional precision—also referred to as increased “transcriptional noise”—observable in different tissues^38–44^.

**Fig. 1.**
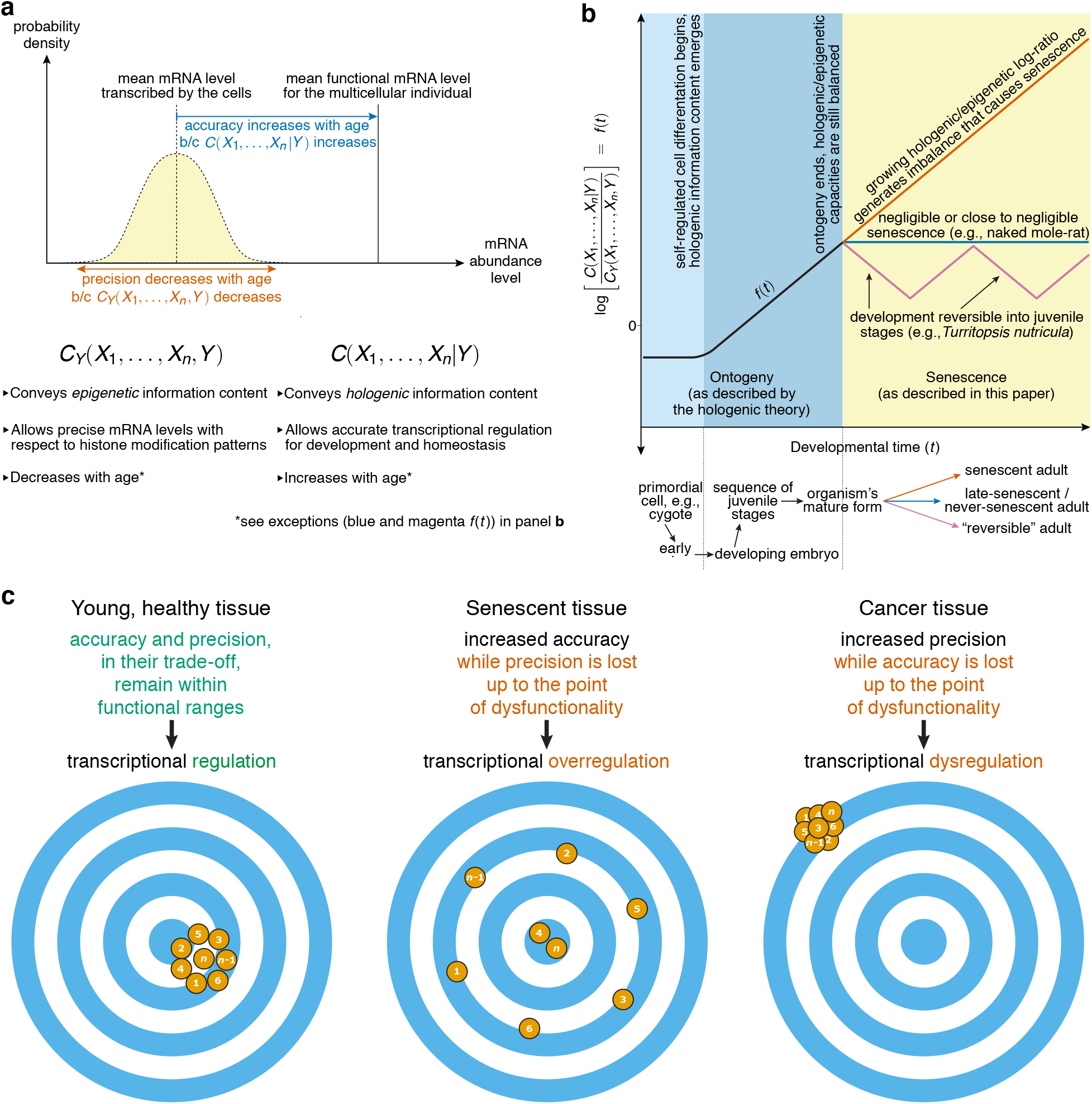
Schematic of the imbalance between information capacity for hologenic content and epigenetic content—and its associated transcriptional accuracy/precision trade-off—in TSS-adjacent histone modification crosstalk as the primary cause of senescence. (a) The age-correlated increase in *C*(*X*_1_,…, *X_n_*|*Y*) and concurrent decrease in *C_Y_*(*X*_1_,…, *X_n_*, *Y*) overregulates the transcription of any given gene with respect to the multicellular individual’s needs, i.e., the transcription of the gene becomes more and more accurate (blue) but less and less precise (orange) up to the point of dysfunctionality. This unavoidable trade-off is explained by histone modification patterns becoming more constrained by regulation at the multicellular-individual level while at the same time becoming worse predictors of mRNA levels. The specific critical level of histone modification crosstalk (i.e., the value of *n* in {*X*_1_,…, *X_n_*}) at which this phenomenon occurs may vary across taxa. (b) The log-ratio of non-epigenetic to epigenetic histone crosstalk magnitude increases during development as the embryo grows (black *f*(*t*) in darker blue area). After the organism reaches its mature form (yellow area), the log-ratio continues to increase (orange *f*(*t*))—with a few notable exceptions (blue and magenta *f*(*t*)). This continuous increase in turn creates an increasing dysfunctional imbalance of information contents that translates into senescence and, eventually, into death. (c) Difference between transcriptional regulation (in young, healthy tissue), overregulation (in senescence), and dysregulation (in cancer) according to this theory, schematized by the transcriptional levels for any given gene in *n* cells (orange dots) of the same type, with the respective bullseye representing the optimal transcriptional level (for the multicellular individual) of said gene in any given condition.

Thus, we can regard senescence under this theory as a global transcriptional overregulation with respect to the multicellular individual’s needs—as opposed to the group of diseases we call cancer, where the dysfunctional dynamics are typically characterized in terms of a dysregulation of transcription and gene expression^45,46^ (see Fig. 1c). Importantly, this global transcriptional overregulation is to be understood under this theory as senescence itself and not as its fundamental cause, which is the previously described epigenetic/non-epigenetic information imbalance in histone crosstalk.

### 2.3. Senescence’s proposed proximate cause

The proximate cause of senescence—i.e., how senescence is triggered throughout adulthood once a multicellular individual reaches its mature form—according to the theory presented here is described in the following steps:

- the histone H1 protein constrains the accessibility of critical histone-modifying enzymes and chromatin remodeling factors to the nucleosome^47,48^
- the histone H1 variants most abundant in somatic, terminally differentiated cells are H1.0 (H1 histone family, member 0; also known as H1°, H1(0), H5, H1δ, or RI H1) and, in vertebrates, also H1x (H1 histone family, member X; also known as H1.10)^49,50^
- variants H1.0 and H1x would undergo post-translational modifications in at least one DNA-binding, nucleosome-proximal amino acid residue within each protein’s globular domain
- these PTMs to H1.0 and H1x histones—such as Ser/Thr phosphorylation, Lys acetylation, and Asn/Gln deamidation—would accumulate, in net terms, throughout adulthood
- the immediate effect of these PTMs would be a decrease in the net electric charge at physiological pH of the DNA-binding (nucleosome-proximal) sites, i.e., rendering their net electric charge less positive, in turn diminishing the DNA-binding affinity in these sites
- in most species this decrease in net electric charge would be quite drastic—potentially even resulting in a locally negative net electric charge—making the residence time of H1.0 and H1x histones in their respective binding site significantly shorter.
- this progressively shorter residence time would underpin the imbalance of constraints described previously, with the core nucleosomal histone (H2A, H2B, H3, and H4) modifications becoming unable to modulate transcriptional levels as they usually do. In other words, H2A, H2B, H3, and H4 PTMs become worse and worse predictors of transcriptional levels—in non-senescent tissue their predictive power is very high (*R* ≈ 0.9)^33^. Importantly, this is a chromatin-wide dysfunctional phenomenon.
- transcription then becomes less and less precise—or “noisier”—up to the point of dysfunctionality
- under the theory presented here, this dysfunctional and ever-increasing loss of transcriptional precision—or, equivalently, gain of “transcriptional noise”—is to be understood as senescence itself

### 2.4. Senescence’s proposed ultimate cause

The theory presented in this paper proposes an alternative ultimate cause. Senescence at the multicellular-individual level is, I suggest, not the result of relaxed selection but instead an intrinsic developmental byproduct that would have been already observable theoretically in the emergence of the very first individuated multicellular organisms as described by the hologenic theory^26^. In other words, had the first individuated multicellular organisms been free from any extrinsic hazard in the wild, they would have begun to senesce significantly after reaching a mature form in their development, as opposed to displaying extremely slow or negligible senescence as can be inferred from the relaxed-selection ultimate cause hypothesis.

## 3. DISCUSSION

### 3.1. Disposable soma or bloated soma?

The theory of senescence presented here may be regarded as a “bloated soma” theory, in the sense that it describes senescence as a dysfunctional byproduct of developmental dynamics, which are not only useful but indeed crucial in early age.

### 3.2. Age-related cancer as a “pushback” against senescence

The delicate balance between hologenic and epigenetic information described here may shed light on the well-known positive correlation between cancer incidence and age^51^: if the senescent multicellular individual attempts to correct its growing hologenic/epigenetic content imbalance too strongly, it may elicit the onset of cancer. Thus, age-related cancer would be the result of a poorly tuned yet strong enough “pushback” from the multicellular individual against its own senescence. Although the specific dynamics that would underpin the “pushback” are beyond the scope of this paper, this hypothesis is indeed falsifiable by means of the following secondary prediction: the observed log-ratio of non-epigenetic to epigenetic histone crosstalk magnitude in the normal (i.e., non-cancerous) cells closest to an age-related stage I malignant tumor will be significantly lower than said log-ratio observed in the other (i.e., tumor-nonadjacent) normal cells of the same tissue. (Note: The falsification of this secondary prediction does not imply the falsification of the theory as a whole.)

In turn, the “pushback”-against-senescence hypothesis for age-related cancer has, if correct, an implication we should not overlook. Namely, stopping senescence efficiently and eliminating the incidence of age-related cancer should be one and the same technical challenge. In this respect, it is worth noting that in the naked mole-rat both senescence^17^ and cancer incidence^52,53^ have been described as negligible or close to negligible.

Rozhok and DeGregori have highlighted the explanatory limitations^54^ of the Armitage-Doll multistage model of carcinogenesis, which regards the accumulation of genetic mutations as the cause of age-related cancer^55^. They further argued that age-related cancer should rather be understood as a function of senescence-related processes^54^. However, their description of age-related cancer is based on Darwinian processes and thus differ from the account suggested here, which can be understood within the concept of teleodynamics^56,57^—a framework of biological individuality based on the emergence of intrinsic higher-order constraints, such as that described in the hologenic theory^26^.

### 3.3. Falsifiability

Falsifiability will be met by the following experimentally testable predictions, which are derived directly from the theory of senescence presented here:

A. For the human histone H1.0 or its most abundant protein orthologs in somatic, terminally differentiated cells of any individuated multicellular species, if its residence time at the binding site is measured—typically as the half-time for recovery, *t*_50_, in a fluorescent recovery after photobleaching (FRAP) assay^58^—with respect to the age of the individual, the observed average slope 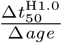 will be non-positive:

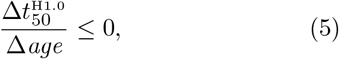

and, for species from the same phylum, also positively correlated with the average adult lifespan of the species, *l*, once log-transformed and controlling for body mass:

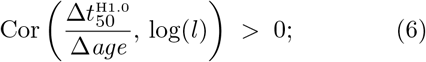

thus, for the shortest-lived species the observed average slope 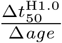 will be steepest (i.e., negative with the highest absolute value) and gentlest (i.e., close or equal to zero) for species displaying negligible senescence.

(Note: When testing this prediction, primary somatic cells should be collected from the same cell type or tissue type.)

B. In any individuated multicellular species and with the random variables *X*_1_,…, *X_n_* representing *n* histone modification levels in specific genomic positions with respect to the TSS and the random variable *Y* representing the per-TSS mRNA abundance in a given (fixed) non-cancerous cell type, there will be a specific level of scale *n*\* such that *C*(*X*_1_,…, *X_n*_*) (i.e., the overall histone crosstalk) will not vary significantly with age:

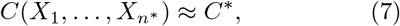

where *C*\* is a constant while, at the same level *n*\*, the following correlation will be both positive and statistically significant:

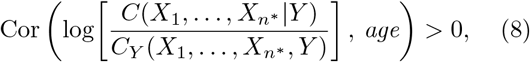

where *age* is that of the indivudual where the respective primary cells for each tandem ChIP-seq and RNA-seq assay were obtained from. Moreover, since hologenic information content is described as emerging locally and independently in each developmental process^26^, the statistical strength of this predicted positive correlation will be further increased—and underpinned by a monotonically increasing function—if all primary cell samples are obtained from the same tissue of the same individual throughout its adulthood (see schematic for testing this prediction in Fig. **2** and further details in Supplementary File 1). The notable exceptions to be made for prediction B are a few species able to undergo reverse developmental processes from adult to juvenile stages. One such species is the jellyfish *Turritopsis nutricula*^59^, which is predicted to display an analogous negative correlation in the processes, i.e., “reassignment” in reverse. Another exception for the predictions are species displaying extremely slow or potentially negligible senescence processes^60^. Examples of these are the bristlecone pine *Pinus longaeva*^61^, the freshwater polyp *Hydra vulgaris*^62^, and the naked mole-rat *Heterocephalus glaber*^17^, which, after adulthood, are predicted to display a significant but very weak positive correlation (in cases where senescence is extremely slow), or an hologenic/epigenetic log-ratio invariant with age (i.e., no correlation in cases where senescence is truly negligible; see Fig. 1b).

**Fig. 2.**
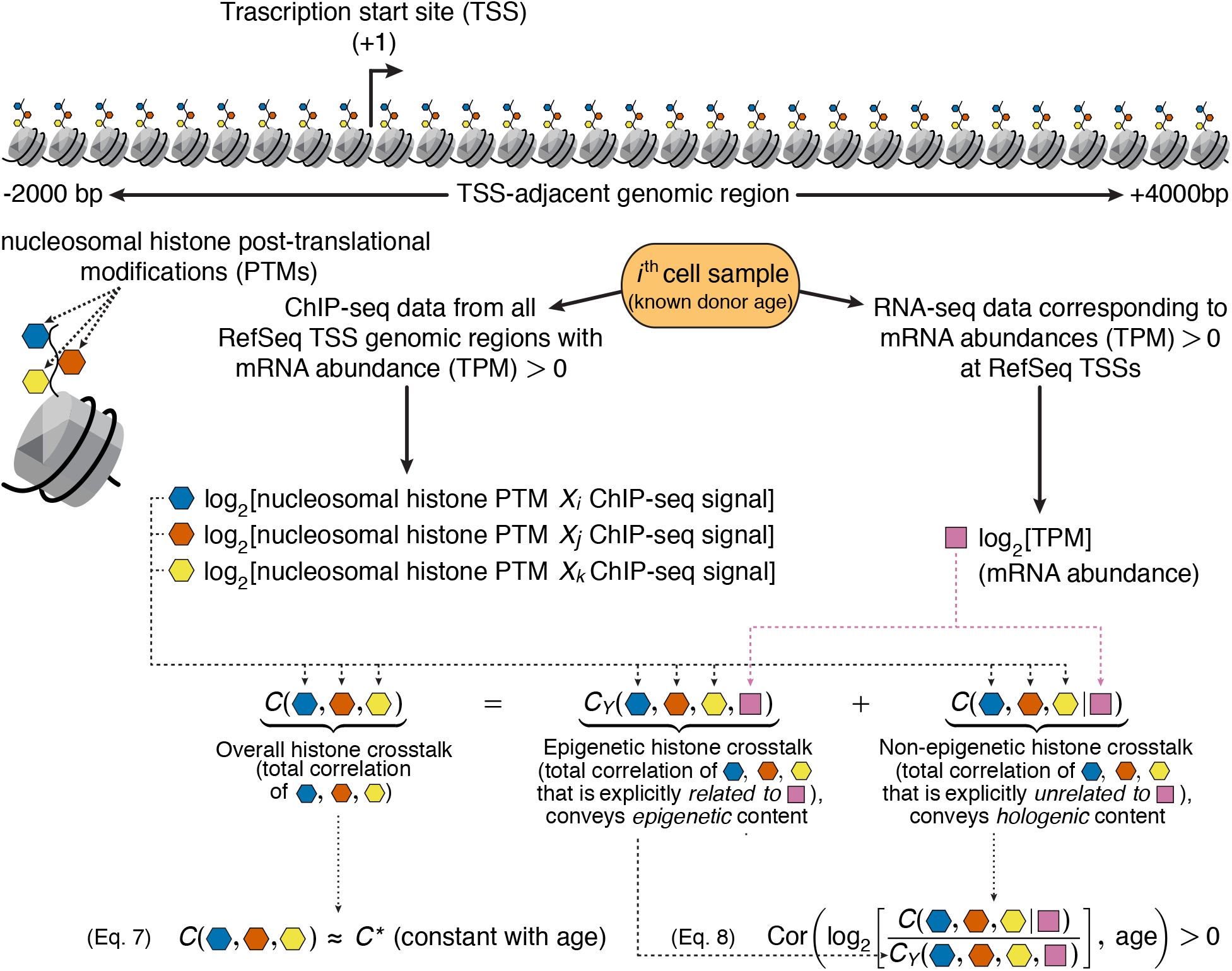
Schematic for testing prediction B using tandem ChIP-seq/RNA-seq from primary cell samples with known donor age. In this case total correlation *C* is estimated using triads {*X_i_*, *X_j_*, *X_k_*} of position-specific ChIP-seq signals with and without respect to the RNA-seq signal *Y*. See details in Supplementary File 1

While predictions A and B can be used to falsify the theory presented here, their verification does not directly support the causal structure proposed in subsection 2.3. In other words, any experimental result verifying said predictions could always be interpreted as being just one of the many effects of senescence—or lack thereof in some species. This is why the following two additional predictions will be provided:

C. If the gene(s) encoding the protein ortholog(s) of the human histone H1.0 most abundant in somatic, terminally differentiated cells of a non-human multicellular species is (are) edited so that (i) a minimum amount (to be determined but predicted to exist) of nucleosome-proximal amino acid residues in the globular domain is substituted by residues that significantly increase the nucleosome-proximal net electric charge, (ii) thereby also significantly increasing the mean residence time of the histone H1.0 ortholog at the DNA binding site, then the mutant population obtained will display a significantly lower mortality rate as a function of age compared to that of a wild-type population. Additionally, if the wild-type species is cancer prone (e.g., mouse), the respective mutant population will be significantly less cancer prone in comparison—see rationale in subsection 3.2. (Note: The nucleosome-proximal amino acid residues in the histone H1.0 globular domain have been already identified^63^ and some of them have been subjected to systematic mutagenesis to study its effect on residence time^58^.)
D. If the gene(s) encoding the protein ortholog(s) of the human histone H1.0 most abundant in somatic, terminally differentiated cells of a non-human multicellular species is (are) edited so that (i) a minimum amount (to be determined but predicted to exist) of nucleosome-proximal amino acid residues in the globular domain is substituted by residues that significantly decrease the nucleosome-proximal net electric charge, (ii) thereby also significantly decreasing the mean residence time of the histone H1.0 ortholog at the DNA binding site, then the mutant population obtained will display a significantly higher mortality rate as a function of age compared to that of a wild-type population. Additionally, if the wild-type species is cancer resistant (e.g., naked mole-rat), the respective mutant population will be significantly less cancer resistant in comparison—see rationale in subsection 3.2.

Importantly, verification of predictions C and D would effectively support the causal structure proposed in subsection 2.3 by providing extremely specific sufficient conditions for directly decreasing or increasing lifespan in any multicellular species.

### 3.4 Concluding remarks

In an upcoming theoretical paper I will specifically address how DNA-binding affinity changes—or lack thereof—in the histone H1.0/H1x proteins when subject to PTMs throughout adulthood may contribute to explain lifespan differences in any multicellular phylum.

## ABBREVIATIONS

H1.0/H1°/H1(0)/H5/H1δ/RI H1: H1 histone family, member 0
H1x/H.10: H1 histone family, member X
PTMs: post-translational modifications
TSSs: transcription start sites

## ACKNOWLEDGEMENTS

I wish to thank Angelika H. Hofmann at SciWri Services for editing this paper into an English I could only hope to write—any departures from that in the current revision are mine and mine alone. I would also like to express my gratitude to Amy Brock for her insightful feedback. I am also indebted to the reviewers for the time they devoted to the evaluation of this paper and their valuable comments.

## Appendix A Shannon measures of statistical uncertainty and statistical association

Shannon measures of statistical uncertainty and statistical association are suggested to be used in order to quantify histone crosstalk at TSSs and its relationship with mRNA levels.

### Statistical uncertainty

C.E. Shannon’s seminal work, among other things, introduced the notion of—and a measure for—the uncertainty about discrete random variables^64^. For a discrete random variable *X* with probability mass function *P*(*X*) its uncertainty (also known as Shannon entropy) is defined as

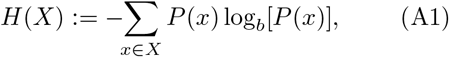

where *P*(*x*) is the probability of *X*=*x* and *b* is the logarithm base. When *b*=2, the unit for this measure is the bit. *H*(*X*) can also be interpreted as the amount of information necessary to resolve the uncertainty about the outcome of *X*. Shannon uncertainty was the measure used to estimate the uncertainty about the mRNA abundance level to be resolved in normal cells.

*H*(*X*) is typically called marginal uncertainty because it involves only one random variable. In a multivariate scenario, the measure *H*(*X*_1_,…, *X_n_*) is called the joint uncertainty of the set of discrete random variables {*X*_1_,…, *X_n_*}, and it is analogously defined as

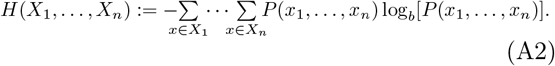

Another measure important to this work is the conditional uncertainty about a discrete random variable *Y*, with probability mass function *P*(*Y*), given that the value of another discrete random variable *X* is known. This conditional uncertainty *H*(*Y*|*X*) can be expressed as

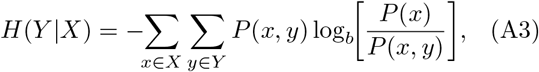

where *P*(*x, y*) is the joint probability of *X*=*x* and *Y*=*y*. Importantly, any measure of Shannon uncertainty (or any other derived Shannon measure) that is conditional on a random variable *X* can also be understood as said measure being explicitly unrelated to, or statistically independent from, the variable *X*.

### Statistical association

A classic Shannon measure of statistical association of any two discrete random variables *X* and *Y* is that of mutual information *I*, defined as

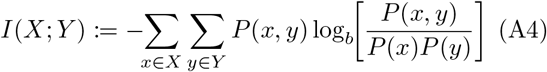

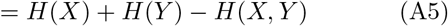

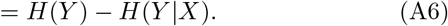

Note that if and only if *X* and *Y* are statistically independent then *I*(*X*; *Y*)=0, *H*(*X, Y*)=*H*(*X*)+*H*(*Y*), and *H*(*Y*|*X*)=*H*(*Y*). To analyze the magnitude of histone H3 crosstalk at TSSs, the two best known multivariate generalizations of mutual information were used in this work. The first is interaction information^65^ or co-information^66^, also symbolized by *I*, which is defined analogously to Eq. A2 for a set *V* of *n* discrete random variables as

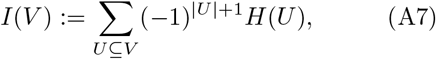

where |*U*| is the cardinality (in this case, the number of random variables) of the subset *U*. In the case of interaction information *I*, Shannon uncertainty *H* is thus summed over all subsets of *V* (the uncertainty of the empty subset is 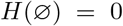). Importantly, the interaction information of the random variables {*X*_1_,…, *X_n_*} can be decomposed with respect to another random variable *Y* as follows:

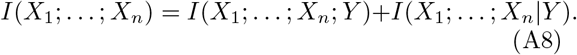

Interaction information *I*(*X*_1_;…; *X_n_*) captures the statistical association of all variables {*X*_1_,…, *X_n_*} taken at once, i.e., excluding all lower-order associations, and it can also take negative values in some cases. Interaction information was used in this work as a means to compute total correlation values.

To specifically quantify the magnitude of histone crosstalk, the second multivariate generalization of mutual information I suggest to be used is total correlation^34^ (symbolized by *C*) or multiinformation^35^, which is defined as

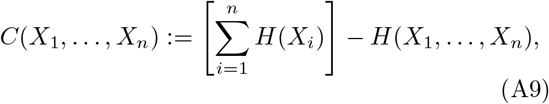

i.e., as the sum of the marginal uncertainties of the random variables {*X*_1_,…, *X_n_*} minus their joint uncertainty. Importantly, and unlike interaction information *I*, total correlation *C* captures all possible statistical associations including lower-order associations or, equivalently, all possible associations between any two or more random variables in the set {*X*_1_,…, *X_n_*}. This is because the definition of interaction information *I* in Eq. A7 allows total correlation *C* to be rewritten as a sum of quantities *I* for all possible combinations of variables in {*X*_1_,…, *X_n_*}:

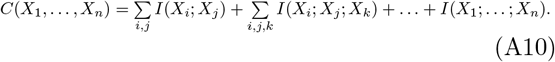

This expression for total correlation *C* as a sum of interaction information quantities *I* along with the sum decomposition of *I* in Eq. A8 allows *C* to be decomposed also as a sum:

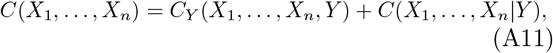

where *C_Y_*(*X*_1_,…, *X_n_, Y*) is the sum (analogous to that of Eq. A10) of all interaction information quantities *I* but now including the random variable *Y* in each combination of variables in {*X*_1_,…, *X_n_*}, i.e.,

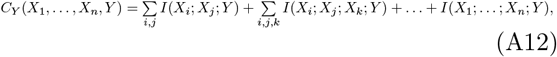

and where *C*(*X*_1_,…, *X_n_*|*Y*) is the sum of all conditional interaction information quantities *I* given *Y* for each combination of variables in {*X*_1_,…, *X_n_*}, i.e.,

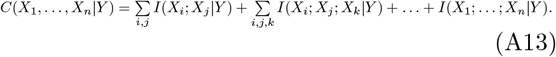

For this work’s purposes, total correlation *C* is proposed as an appropriate measure of statistical association to assess TSS-adjacent histone crosstalk because (i) *C* is non-negative and thus easier to interpret conceptually, (ii) *C* is equal to zero if and only if all random variables it comprises are statistically independent, (iii) *C* captures all possible associations up to a given number of variables (in this work, position-specific histone modification levels) and, (iv) *C* can be decomposed, as shown in Eq. A11, as a sum of two *C* quantities: one explicitly related to a certain variable *Y* and the other explicitly unrelated to *Y*. Property (iv) was useful to decompose the overall histone crosstalk as a sum of an epigenetic and a non-epigenetic component (Eq. 1).

An additional Shannon measure of statistical association can be used to assess the predictive power of TSS-adjacent histone modification levels on mRNA abundance levels (such power has already been used to predict mRNA levels with high accuracy^33^). The uncertainty coefficient *U*^67^ is defined as

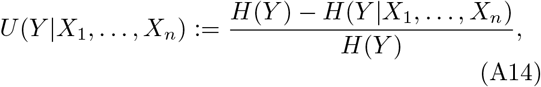

i.e., *U*(*Y*|*X*_1_,…, *X_n_*) is the relative decrease in uncertainty about *Y* when {*X*_1_,…, *X_n_*} are known—or, equivalently, the fraction of bits in *Y* that can be predicted by {*X*_1_,…, *X_n_*}—and it can take values from 0 to 1. *U*(*Y*|*X*_1_,…, *X_n_*)=0 implies the set {*X*_1_,…, *X*_n_} has no predictive power on *Y*, whereas *U*(*Y*|*X*_1_,…, *X_n_*)=1 implies {*X*_1_,…, *X_n_*} can predict *Y* completely.

## REFERENCES

1 S. Katz, Imagining the life-span: From premodern miracles to postmodern fantasies, in: Images of Aging: Cultural Representations of Later Life, Routledge, London, 1995, pp. 59–74.

2 J. Vijg, J. Campisi, Puzzles, promises and a cure for ageing, Nature 454 (2008) 1065–1071. doi:10.1038/nature07216.

3 J. W. Shay, W. E. Wright, Hayflick, his limit, and cellular ageing, Nat. Rev. Mol. Cell Biol. 1 (2000) 72–76. doi:10.1038/35036093.

4 L. Hayflick, P. S. Moorhead, The serial cultivation of human diploid cell strains, Exp. Cell Res. 25 (3) (1961) 585–621. doi:10.1016/0014-4827(61)90192-6.

5 L. Roger, F. Tomas, V. Gire, Mechanisms and regulation of cellular senescence, Int. J. Mol. Sci. 22 (23) (2021) 1–42. doi:10.3390/ijms222313173.

6 V. J. Cristofalo, R. G. Allen, R. J. Pignolo, B. G. Martin, J. C. Beck, Relationship between donor age and the replicative lifespan of human cells in culture: A reevaluation, Proc. Natl. Acad. Sci. 95 (18) (1998) 10614–10619. URL https://www.pnas.org/content/95/18/10614

7 A. Lorenzini, M. Tresini, S. N. Austad, V. J. Cristofalo, Cellular replicative capacity correlates primarily with species body mass not longevity, Mech. Ageing Dev. 126 (10) (2005) 1130–1133. doi:10.1016/j.mad.2005.05.004.

8 C. López-Otín, M. A. Blasco, L. Partridge, M. Serrano, G. Kroemer, The hallmarks of aging, Cell 153 (6) (2013) 1194–1217. doi:10.1016/j.cell.2013.05.039.

9 E. Mayr, Cause and effect in biology, Science 134 (3489) (1961) 1501–1506. doi:10.1126/science.134.3489.1501.

10 J. B. S. Haldane, New paths in genetics, George Allen & Unwin, London, 1941.

11 P. B. Medawar, An unsolved problem of biology, H. K. Lewis, London, 1952.

12 G. C. Williams, Pleiotropy, natural selection, and the evolution of senescence, Evolution 11 (4) (1957) 398–411. doi:10.2307/2406060.

13 T. B. L. Kirkwood, Evolution of ageing, Nature 270 (1977) 301–304. doi:10.1038/270301a0.

14 T. B. Kirkwood, Understanding the odd science of aging, Cell 120 (4) (2005) 437–447. doi:10.1016/j.cell.2005.01.027.

15 T. Flatt, Survival costs of reproduction in *Drosophila*, Exp. Gerontol. 46 (5) (2011) 369–375. doi:10.1016/j.exger.2010.10.008.

16 R. Buffenstein, The naked mole-rat: a new long-living model for human aging research, Journals Gerontol. Ser. A Biol. Sci. Med. Sci. 60 (11) (2005) 1369–1377. doi:10.1093/gerona/60.11.1369.

17 J. G. Ruby, M. Smith, R. Buffenstein, Naked mole-rat mortality rates defy Gompertzian laws by not increasing with age, Elife 7 (2018) e31157. doi:10.7554/eLife.31157.

18 K. Jin, Modern biological theories of aging, Aging Dis. 1 (2) (2010) 72–74. URL http://www.aginganddisease.org/EN/Y2010/V1/I2/72

19 M. V. Blagosklonny, Aging is not programmed: Genetic pseudo-program is a shadow of developmental growth, Cell Cycle 12 (24) (2013) 3736–3742. doi:10.4161/cc.27188.

20 S. Horvath, K. Raj, DNA methylation-based biomarkers and the epigenetic clock theory of ageing, Nat. Rev. Genet. doi:10.1038/s41576-018-0004-3.

21 R. W. Powers 3rd, M. Kaeberlein, S. D. Caldwell, B. K. Kennedy, S. Fields, Extension of chronological life span in yeast by decreased TOR pathway signaling, Genes Dev. 20 (2) (2006) 174–184. doi:10.1101/gad.1381406.

22 C. Kenyon, J. Chang, E. Gensch, A. Rudner, R. Tabtiang, A *C. elegans* mutant that lives twice as long as wild type, Nature 366 (1993) 461–464. doi:10.1038/366461a0.

23 T. Vellai, K. Takacs-Vellai, Y. Zhang, A. L. Kovacs, L. Orosz, F. Müller, Genetics: Influence of TOR kinase on lifespan in *C. elegans*, Nature 426 (2003) 620. doi:10.1038/426620a.

24 D. Gems, L. Partridge, Genetics of longevity in model organisms: debates and paradigm shifts, Annu. Rev. Physiol. 75 (1) (2013) 621–644. doi:10.1146/annurev-physiol-030212-183712.

25 M. Yousefzadeh, C. Henpita, R. Vyas, C. Soto-Palma, P. Robbins, L. Niedernhofer, DNA damage—how and why we age?, Elife 10 (2021) e62852. doi:10.7554/eLife.62852.

26 F. A. Veloso, On the developmental self-regulatory dynamics and evolution of individuated multicellular organisms, J. Theor. Biol. 417 (2017) 84–99. doi:10.1016/j.jtbi.2016.12.025.

27 P. W. Atkins, The Second Law, Scientific American Library, New York, 1984.

28 J.-S. Lee, E. Smith, A. Shilatifard, The language of histone crosstalk, Cell 142 (5) (2010) 682–685. doi:10.1016/j.cell.2010.08.011.

29 Z. Wang, C. Zang, J. A. Rosenfeld, D. E. Schones, A. Barski, S. Cuddapah, K. Cui, T.-Y. Roh, W. Peng, M. Q. Zhang, K. Zhao, Combinatorial patterns of histone acetylations and methylations in the human genome, Nat Genet. 40 (7) (2008) 897–903. doi:10.1038/ng.154.

30 A. Lennartsson, K. Ekwall, Histone modification patterns and epigenetic codes, Biochim. Biophys. Acta, 1790 (9) (2009) 863–868. doi:10.1016/j.bbagen.2008.12.006.

31 V. E. A. Russo, R. A. Martienssen, A. D. Riggs, Epigenetic mechanisms of gene regulation, Cold Spring Harbor Laboratory Press, Cold Spring Harbor, 1996.

32 Y. Wang, Q. Yuan, L. Xie, Histone modifications in aging: the underlying mechanisms and implications, Curr. Stem Cell Res. Ther. 13 (2) (2018) 125–135. doi:10.2174/1574888x12666170817141921.

33 V. Kumar, M. Muratani, N. A. Rayan, P. Kraus, T. Lufkin, H. H. Ng, S. Prabhakar, Uniform, optimal signal processing of mapped deep-sequencing data, Nat. Biotechnol. 31 (7) (2013) 615–622. doi:10.1038/nbt.2596.

34 S. Watanabe, Information theoretical analysis of multivariate correlation, IBM J. Res. Dev. 4 (1) (1960) 66–82. doi:10.1147/rd.41.0066.

35 M. Studený, J. Vejnarová, The multiinformation function as a tool for measuring stochastic dependence, in: Learning in Graphical Models, 1998, pp. 261–297. doi:10.1007/978-94-011-5014-9\_10.

36 G. Csárdi, A. Franks, D. S. Choi, E. M. Airoldi, D. A. Drummond, Accounting for experimental noise reveals that mRNA levels, amplified by post-transcriptional processes, largely determine steady-state protein levels in yeast, PLoS Genet. 11 (5) (2015) e1005206. doi:10.1371/journal.pgen.1005206.

37 S. E. Prochnik, J. Umen, A. M. Nedelcu, A. Hallmann, S. M. Miller, I. Nishii, P. Ferris, A. Kuo, T. Mitros, L. K. Fritz-Laylin, U. Hellsten, J. Chapman, O. Simakov, S. A. Rensing, A. Terry, J. Pangilinan, V. Kapitonov, J. Jurka, A. Salamov, H. Shapiro, J. Schmutz, J. Grimwood, E. Lindquist, S. Lucas, I. V. Grigoriev, R. Schmitt, D. Kirk, D. S. Rokhsar, Genomic analysis of organismal complexity in the multicellular green alga *Volvox carteri*, Science 329 (5988) (2010) 223–226. doi:10.1126/science.1188800.

38 M. C. Salzer, A. Lafzi, A. Berenguer-Llergo, C. Youssif, A. Castellanos, G. Solanas, F. O. Peixoto, C. Stephan-Otto Attolini, N. Prats, M. Aguilera, J. Martín-Caballero, H. Heyn, S. A. Benitah, Identity noise and adipogenic traits characterize dermal fibroblast aging, Cell 175 (6) (2018) 1575–1590.e22. doi:10.1016/j.cell.2018.10.012.

39 R. Bahar, C. H. Hartmann, K. A. Rodriguez, A. D. Denny, R. A. Busuttil, M. E. Dollé, R. B. Calder, G. B. Chisholm, B. H. Pollock, C. A. Klein, J. Vijg, Increased cell-to-cell variation in gene expression in ageing mouse heart, Nature 441 (2006) 1011–1014. doi:10.1038/nature04844.

40 I. Angelidis, L. M. Simon, I. E. Fernandez, M. Strunz, C. H. Mayr, F. R. Greiffo, G. Tsitsiridis, M. Ansari, E. Graf, T. M. Strom, M. Nagendran, T. Desai, O. Eickelberg, M. Mann, F. J. Theis, H. B. Schiller, An atlas of the aging lung mapped by single cell transcriptomics and deep tissue proteomics, Nat. Commun. 10 (963) (2019). doi:10.1038/s41467-019-08831-9.

41 C. Nikopoulou, S. Parekh, P. Tessarz, Ageing and sources of transcriptional heterogeneity, Biol. Chem. 400 (7) (2019) 867–878. doi:10.1515/hsz-2018-0449.

42 C. P. Martinez-Jimenez, N. Eling, H.-C. Chen, C. A. Vallejos, A. A. Kolodziejczyk, F. Connor, L. Stojic, T. F. Rayner, M. J. T. Stubbington, S. A. Teichmann, Others, Aging increases cell-to-cell transcriptional variability upon immune stimulation, Science 355 (6332) (2017) 1433–1436. doi:10.1126/science.aah4115.

43 M. Enge, H. E. Arda, M. Mignardi, J. Beausang, R. Bottino, S. K. Kim, S. R. Quake, Single-cell analysis of human pancreas reveals transcriptional signatures of aging and somatic mutation patterns, Cell 171 (2) (2017) 321–330.e14. doi:10.1016/j.cell.2017.09.004.

44 A. Swisa, K. H. Kaestner, Y. Dor, Transcriptional noise and somatic mutations in the aging pancreas, Cell Metab. 26 (6) (2017) 809–811. doi:10.1016/j.cmet.2017.11.009.

45 K. Malik, K. W. Brown, Epigenetic gene deregulation in cancer, Br. J. Cancer 83 (2000) 1583–1588. doi:10.1054/bjoc.2000.1549.

46 T. J. Gonda, R. G. Ramsay, Directly targeting transcriptional dysregulation in cancer, Nat. Rev. Cancer 15 (2015) 686–694. doi:10.1038/nrc4018.

47 N. Happel, D. Doenecke, Histone H1 and its isoforms: contribution to chromatin structure and function, Gene 431 (1-2) (2009) 1–12. doi:10.1016/j.gene.2008.11.003.

48 F. Song, P. Chen, D. Sun, M. Wang, L. Dong, D. Liang, R.-M. Xu, P. Zhu, G. Li, Cryo-EM study of the chromatin fiber reveals a double helix twisted by tetranucleosomal units, Science 344 (6182) (2014) 376–380. doi:10.1126/science.1251413.

49 P. B. Talbert, K. Ahmad, G. Almouzni, J. Ausió, F. Berger, P. L. Bhalla, W. M. Bonner, W. Z. Cande, B. P. Chadwick, S. W. L. Chan, Others, A unified phylogeny-based nomenclature for histone variants, Epigenetics & Chromatin 7 (5) (2012). doi:10.1186/1756-8935-5-7.

50 E. J. Draizen, A. K. Shaytan, L. Mariño-Ramírez, P. B. Talbert, D. Landsman, A. R. Panchenko, HistoneDB 2.0: a histone database with variants—an integrated resource to explore histones and their variants, Database 2016 (2016) baw014. doi:10.1093/database/baw014.

51 F. Kamangar, G. M. Dores, W. F. Anderson, Patterns of cancer incidence, mortality, and prevalence across five continents: defining priorities to reduce cancer disparities in different geographic regions of the world, J. Clin. Oncol. 24 (14) (2006) 2137–2150. doi:10.1200/JCO.2005.05.2308.

52 R. Buffenstein, Negligible senescence in the longest living rodent, the naked mole-rat: Insights from a successfully aging species, J. Comp. Physiol. B Biochem. Syst. Environ. Physiol. 178 (4) (2008) 439–445. doi:10.1007/s00360-007-0237-5.

53 M. A. Delaney, L. Nagy, M. J. Kinsel, P. M. Treuting, Spontaneous histologic lesions of the adult naked mole rat (*Heterocephalus glaber*): A retrospective survey of lesions in a zoo population, Vet. Pathol. 50 (4) (2013) 607–621. doi:10.1177/0300985812471543.

54 A. I. Rozhok, J. DeGregori, The evolution of lifespan and age-dependent cancer risk, Trends in Cancer 2 (10) (2016) 552–560. doi:10.1016/j.trecan.2016.09.004.

55 P. Armitage, R. Doll, The age distribution of cancer and a multi-stage theory of carcinogenesis, Br. J. Cancer 8 (1954) 1–12. doi:10.1038/bjc.1954.1.

56 T. W. Deacon, Incomplete Nature: How Mind Emerged from Matter, W.W. Norton & Company, New York, 2011.

57 T. W. Deacon, A. Srivastava, J. A. Bacigalupi, The transition from constraint to regulation at the origin of life, Front. Biosci. 19 (2014) 945–957. doi:10.2741/4259.

58 D. T. Brown, T. Izard, T. Misteli, Mapping the interaction surface of linker histone H10 with the nucleosome of native chromatin in vivo, Nat. Struct. Mol. Biol. 13 (3) (2006) 250–255. doi:10.1038/nsmb1050.

59 S. Piraino, F. Boero, B. Aeschbach, V. Schmid, Reversing the life cycle: Medusae transforming into polyps and cell transdifferentiation in *Turritopsis nutricula* (Cnidaria, Hydrozoa), Biol. Bull. 190 (3) (1996) 302–312. doi:10.2307/1543022.

60 C. E. Finch, Update on slow aging and negligible senescence—a mini-review, Gerontology 55 (3) (2009) 307–313. doi:10.1159/000215589.

61 R. M. Lanner, K. F. Connor, Does bristlecone pine senesce?, Exp. Gerontol. 36 (4-6) (2001) 675–685. doi:10.1016/S0531-5565(00)00234-5.

62 R. Schaible, A. Scheuerlein, M. J. Dańko, J. Gampe, D. E. Martínez, J. W. Vaupel, Constant mortality and fertility over age in *Hydra*, Proc. Natl. Acad. Sci. 112 (51) (2015) 15701–15706. doi:10.1073/pnas.1521002112.

63 J. Bednar, I. Garcia-Saez, R. Boopathi, A. R. Cutter, G. Papai, A. Reymer, S. H. Syed, I. N. Lone, O. Tonchev, C. Crucifix, H. Menoni, C. Papin, D. A. Skoufias, H. Kurumizaka, R. Lavery, A. Hamiche, J. J. Hayes, P. Schultz, D. Angelov, C. Petosa, S. Dimitrov, Structure and dynamics of a 197 bp nucleosome in complex with linker histone H1, Mol. Cell 66 (3) (2017) 384–397.e8. doi:10.1016/j.molcel.2017.04.012.

64 C. E. Shannon, A mathematical theory of communication, Bell Syst. Tech. J. 27 (3) (1948) 379–423. doi:10.1002/j.1538-7305.1948.tb01338.x.

65 W. McGill, Multivariate information transmission, Trans. IRE Prof. Gr. Inf. Theory 4 (4) (1954) 93–111. doi:10.1109/tit.1954.1057469.

66 A. J. Bell, The co-information lattice, Proc. 4th Int. Symp. Independent Component Analysis and Blind Source Separation. URL http://citeseerx.ist.psu.edu/viewdoc/summary?doi=10.1.1.320.5264

67 W. H. Press, S. A. Teukolsky, W. T. Vetterling, B. P. Flannery, Numerical Recipes in C: The Art of Scientific Computing, Cambridge Univ. Press, New York, 1992.

68 T. W. Deacon, How molecules became signs, Biosemiotics (2021). doi:10.1007/s12304-021-09453-9.

69 F. A. Veloso, Interpretation as a form of thermodynamic work, Biosemiotics (2021). doi:10.1007/s12304-021-09456-6.

