## Supplementary File 1 for "A constraints-based theory of the primary cause of senescence: imbalance of epigenetic and non-epigenetic information in histone crosstalk"

### A METHOD FOR TESTING PREDICTION B IN TANDEM CHIP-SEQ (HISTONE H3 MODIFICATIONS) AND RNA-SEQ DATA FROM HUMAN PRIMARY CELLS

The method outlined here mirrors that used by Kumar *et al.* [1] to predict mRNA levels from nucleosomal histone modification profiles with high accuracy.

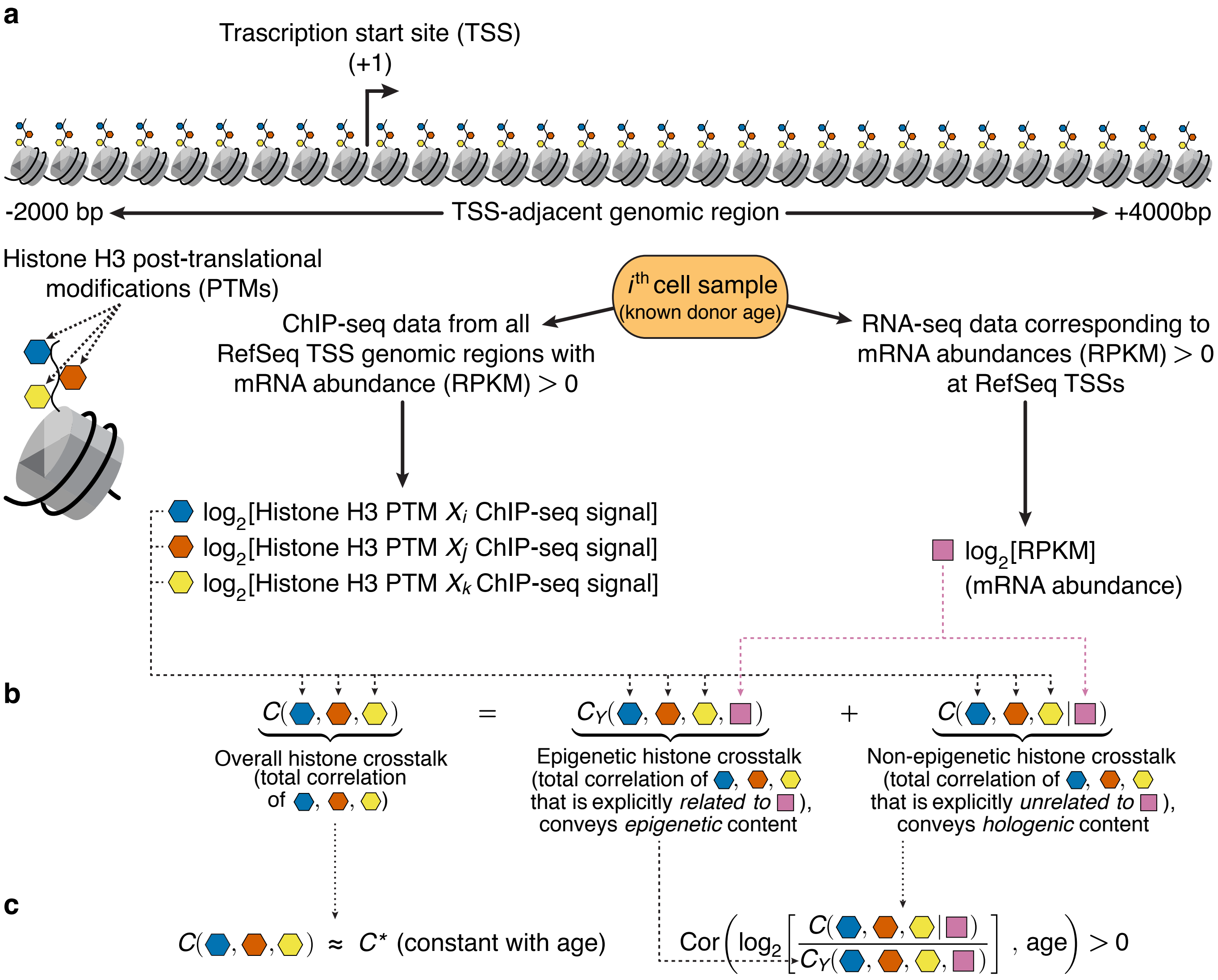


**Fig. 1. Schematic of the computational analysis—similar in method to that used in [1]—for testing prediction B of the theory.** Publicly available, tandem ChIP-seq (chromatin immunoprecipitation followed by high-throughput DNA sequencing) and RNA-seq (transcriptome high-throughput sequencing) data for (e.g., human) primary cell samples allow the computation, for each TSS, of position-specific nucleosomal histone (e.g., histone H3) modification levels (at every 200bp) and its associated mRNA abundance level **(a)**. After log-transforming these levels and taking into account all TSSs, the TSS-adjacent histone H3 crosstalk (triad-wise crosstalk depicted here) can be represented as a total correlation [2] or information capacity in bits, which in turn can be decomposed as the sum of two measurable and explicitly unrelated components: one epigenetic (explicitly related to transcriptional changes) and the other non-epigenetic (explicitly unrelated to said changes) **(b)**. Taking into account all samples, the overall histone crosstalk is predicted to remain statistically constant in time **(c, left)** and log-ratio of non-epigenetic to epigenetic histone H3 crosstalk magnitude is predicted to be positively correlated with cell donor age in normal cells **(c, left)**—and also to be uncorrelated with cell donor age in cancer cells.

#### Data collection

The genomic coordinates and associated transcript lengths of all annotated RefSeq mRNA TSSs for the hg19 (*Homo sapiens*) assembly can be downloaded from the UCSC (University of California, Santa Cruz) database [3]. ChIP-seq and RNA-seq data can be downloaded from the Canadian Epigenetics, Epigenomics, Environment and Health Research Consortium (CEEHRC).

Cell sample data should be selected based on the following criteria: (i) only data sets with associated age are to be included (so prediction B can be tested since involves age) (ii) among these data sets, the group that maximizes the number of specific histone H3 modifications present in all data sets.

#### ChIP-seq datafile processing

Original ChIP-seq binary datafile format is typically bigWig. For mapping the ChIP-seq signal into the hg19 assembly, each datafile can be processed with standard bioinformatics tools [4–6] in the following pipeline:

```
bigWigToWig → wig2bed --zero-indexed → sort -k1,1 -k2,2n →
bedtools map -o median -null 0 -a hg19_all_tss.bed/hg19_all_tss_control.bed
```

to generate an associated BED (Browser Extensible Data) file. (Note: The `hg19_all_tss.bed` file will be a 200bp-per-bin BED reference file with no score values to perform the final ChIP-seq histone modification data mapping onto the 6,000bp-long TSS-adjacent genomic regions. The `hg19_all_tss_control.bed` file will be an analogous BED reference file for mapping the ChIP-seq input data onto 200-bp, 1-kbp, 5-kbp, and 10-kbp genomic windows, see ChIP-seq read profiles and normalization.)

#### ChIP-seq read profiles and normalization

To quantify and represent ChIP-seq read signal profiles for the histone H3 modifications, data can be processed with the same method used in the EFilter multivariate algorithm [1] to predict mRNA levels with high accuracy ($R \approx 0.9$) as mentioned earlier. Steps in said method comprise (i) dividing the genomic region from 2 kbp upstream to 4 kbp downstream of each TSS into 30 200-bp-long bins, in each of which ChIP-seq reads were later counted; (ii) dividing the read count signal for each bin by its corresponding control (ChIP-seq input) read density to minimize artifactual peaks; (iii) estimating the control read density within a 1-kbp window centered on each bin, if the 1-kbp window contained at least 20 reads; otherwise, a 5-kbp window, or else a 10-kbp window was used if the control reads were less than 20. When the 10-kbp length was insufficient, a pseudo-count value of 20 reads per 10 kbp was set as the control read density. This implies that the denominator (i.e., control read density) is at least 0.4 reads per bin.

#### RNA-seq datafile processing

For each strand of DNA, original datafiles typically contain mRNA abundances in RPKM (reads per kilobase of transcript per million mapped reads) in bigWig format. These datafiles can be thus processed analogously to the ChIP-seq datafiles, i.e., using the pipeline

```
bigWigToWig → wig2bed --zero-indexed → sort -k1,1 -k2,2n →
bedtools map -o median -null 0 -a refseq_pos.bed/refseq_neg.bed
```

to obtain associated BED files. (Note: The `refseq_pos.bed` and `refseq_neg.bed` files will be BED reference files for each strand, with no score values, to perform the final RPKM calculation for each RefSeq mRNA in the hg19 assembly.)

If two or more mRNAs share the same TSS (i.e., transcription start site with same genomic position and strand) the mean of the respective RPKM values should be computed and associated with the corresponding TSS.

(Note: In previous work it has been argued that RPKM may not always be a suitable unit of mRNA abundance when studying differential gene expression. Specifically, it was shown that, if transcript size distribution varies significantly among the samples, RPKM might introduce significant biases [7]. To overcome this problem, an alternative abundance unit TPM (transcripts per million)—which is an invertible linear transformation of the RPKM value for each sample—was introduced [7]. Nonetheless, this issue was not a problem for the present work because Shannon measures are invariant under any invertible transformation of the discrete random variables.)

##### ChIP-seq/RNA-seq signal data tables

Using the RPKM values processed in this work, a subset $\mathrm{TSS}_{\mathrm{def}}$ of all RefSeq mRNA TSSs displaying measured abundance (i.e., RPKM $> 0$) in all samples must be identified. The obtained $\mathrm{TSS}_{\mathrm{def}}$ subset will thus provide the data analysis with a common basis for all samples that comprises most protein-coding genes annotated in the human genome.

Following the approach by Kumar *et al.*, for each sample data entry, 30 genomic bins are to be defined and denoted by the distance (bp) between their 5′-end and their respective $\mathrm{TSS}_{\mathrm{def}}$ genomic coordinate: "-2000", "-1800", "-1600", "-1400", "-1200", "-1000", "-800", "-600", "-400", "-200", "0" ($\mathrm{TSS}_{\mathrm{def}}$ or '+1'), "200", "400", "600", "800", "1000", "1200", "1400", "1600", "1800", "2000", "2200", "2400", "2600", "2800", "3000", "3200", "3400", "3600", and "3800". Then, for each sample data entry, the ChIP-seq read signal must be computed for all bins and for all histone modifications (30 bins×5 modifications=150 signal values) in all $\mathrm{TSS}_{\mathrm{def}}$ genomic regions. Data input tables—comprising in this example the histone H3 modifications H3K4me1, H3K9me3, H3K27ac, H3K27me3, and H3K36me3—must thus be generated for each sample entry as:

| | H3K4me1_-2000 | ... | H3K36me3_-2000 | ··· | H3K4me1_3800 | ... | H3K36me3_3800 | RPKM |
|---|---|---|---|---|---|---|---|---|
| #$\mathrm{TSS}_{\mathrm{def}}$ | 4.68 | ... | 7.94 | ··· | 1.32 | ... | 12.15 | 35.63 |
| | ⋮ | | ⋮ | | ⋮ | | ⋮ | ⋮ |
| | 2.13 | ... | 4.97 | ··· | 6.33 | ... | 3.06 | 17.44 |

The tables can then be written into tab-delimited datafiles.

##### Software availability for computing Shannon measures

The `R` software [8] and its `infotheo` package [9] can be used for the computation of Shannon measures of statistical uncertainty and statistical association from the aforementioned datafiles. In particular, marginal and joint Shannon uncertainties and all the other derived Shannon measures may be computed using maximum likelihood (ML) estimation [10] and bias-corrected with the Miller-Madow method [11].

##### Levels of possible statistical associations when assessing histone crosstalk magnitudes

An important aspect of quantifying the epigenetic and non-epigenetic histone crosstalk components is the specific range of possible statistical associations. In other words, the choice of the number $n$ of TSS-adjacent, position-specific histone H3 modification levels when computing $C_Y(X_1,\ldots,X_n,Y)$ and $C(X_1,\ldots,X_n|Y)$. To this end, the minimal $n$ able to predict mRNA levels significantly and non-redundantly—which corresponds to the level of histone crosstalk able to convey a non-neglectable amount of epigenetic information content—must be determined. This value is straightforward to assess using the uncertainty coefficient $U(Y|X_1,\ldots,X_n)$, where $Y$ represents mRNA levels.

In effect, $U(Y|X_i,X_j)$ (i.e., where $n$=2) quantifies the predictive power of pairs of position-specific histone modification levels, $U(Y|X_i,X_j,X_k)$ quantifies the predictive power of triads, etc. $U(Y|X_1,\ldots,X_n)$ values were thus computed for singletons, pairs, triads, and tetrads. Singeltons (i.e., $U(Y|X_i)$) may be calculated for descriptive purposes only, because histone crosstalk is not defined for singletons. In this context, it is useful to assess the synergy of a set of predictor variables [12]. For a triad of variables $\{X_i,X_j,X_k\}$ with respect to a response variable $Y$, synergy is defined as:

$$\frac{U(Y|X_i,X_j,X_k)}{\sum\limits_i U(Y|X_i)+\sum\limits_{i,j} U(Y|X_i,X_j)}, \tag{1}$$

In other words, synergy is the predictive power of a triad divided by the sum of the predictive power of all singletons and pairs within the triad. For a tetrad, similarly, the denominator would have the predictive power of all singletons, pairs, and triads within the tetrad.

It is expected that for all possible singletons ($U(Y|X_i)$), pairs ($U(Y|X_i,X_j)$), triads ($U(Y|X_i,X_j,X_k)$), etc., the predictive power will be increasing, at one point even dramatically (i.e., high synergy; with triads, for example) but if one extra predictor variable is added (to form a tetrad now) and synergy falls dramatically, then the level we are looking for (high predictive power and high variable synergy) will be $n = 3$ (triads).

The example data table has (30 bins×5 modifications=150 predictor variables):

| | H3K4me1_-2000 | ... | H3K36me3_-2000 | ··· | H3K4me1_3800 | ... | H3K36me3_3800 | RPKM |
|---|---|---|---|---|---|---|---|---|
| | 4.68 | ... | 7.94 | ··· | 1.32 | ... | 12.15 | 35.63 |
| #TSS$_{\text{def}}$ | ⋮ | | ⋮ | | ⋮ | | ⋮ | ⋮ |
| | 2.13 | ... | 4.97 | ··· | 6.33 | ... | 3.06 | 17.44 |

In other words, in this example triads $\{X_i, X_j, X_k\}$ would be used out of $\{X_1, \ldots, X_{150}\}$ predictor variables (i.e., columns in the table except by the last column with the transcript abundance values). This implies there are 551,300 possible triads.

In this context, for testing prediction B, 551,300 values of

$$C^* = C(X_i, X_j, X_k) \tag{2}$$

and 551,300 values of

$$r = \mathrm{Cor}\left(\log\left[\frac{C(X_i, X_j, X_k|Y)}{C_Y(X_i, X_j, X_k, Y)}\right],\ age\right) \tag{3}$$

will be computed.

Depending on the empirical distributions obtained for $C^*$ and $r$ (e.g., the distribution of correlation coefficients ($r$) is known to be non-Gaussian [13]), adequate statistical tests and correction for multiple comparisons will have to be used in order to establish a definitive conclusion.

#### REFERENCES

[1] V. Kumar, M. Muratani, N. A. Rayan, P. Kraus, T. Lufkin, H. H. Ng, S. Prabhakar, Uniform, optimal signal processing of mapped deep-sequencing data, Nat. Biotechnol. 31 (7) (2013) 615–622. DOI:10.1038/nbt.2596.

[2] S. Watanabe, Information theoretical analysis of multivariate correlation, IBM J. Res. Dev. 4 (1) (1960) 66–82. DOI:10.1147/rd.41.0066.

[3] D. Karolchik, A. S. Hinrichs, T. S. Furey, K. M. Roskin, C. W. Sugnet, D. Haussler, W. J. Kent, The UCSC Table Browser data retrieval tool, Nucleic Acids Res. 32 (Database issue) (2004) D493–6. DOI:10.1093/nar/gkh103.

[4] W. J. Kent, A. S. Zweig, G. Barber, A. S. Hinrichs, D. Karolchik, BigWig and BigBed: enabling browsing of large distributed datasets, Bioinformatics 26 (17) (2010) 2204–2207. DOI:10.1093/bioinformatics/btq351.

[5] S. Neph, M. S. Kuehn, A. P. Reynolds, E. Haugen, R. E. Thurman, A. K. Johnson, E. Rynes, M. T. Maurano, J. Vierstra, S. Thomas, R. Sandstrom, R. Humbert, J. A. Stamatoyannopoulos, BEDOPS: high-performance genomic feature operations, Bioinformatics 28 (14) (2012) 1919–1920. DOI:10.1093/bioinformatics/bts277.

[6] A. R. Quinlan, I. M. Hall, BEDTools: a flexible suite of utilities for comparing genomic features, Bioinformatics 26 (6) (2010) 841–842. DOI:10.1093/bioinformatics/btq033.

[7] G. P. Wagner, K. Kin, V. J. Lynch, Measurement of mRNA abundance using RNA-seq data: RPKM measure is inconsistent among samples, Theory Biosci. 131 (4) (2012) 281–285. DOI:10.1007/s12064-012-0162-3.

[8] R Core Team, R: a language and environment for statistical computing (2017). Available from: https://www.r-project.org/.

[9] P. E. Meyer, infotheo: information-theoretic measures (2014). Available from: https://cran.r-project.org/package=infotheo.

[10] L. Paninski, Estimation of entropy and mutual information, Neural Comput. 15 (6) (2003) 1191–1253. DOI:10.1162/089976603321780272.

[11] G. A. Miller, Note on the bias of information estimates, Inf. theory Psychol. Probl. methods 2 (95) (1955) 100.

[12] D. Chicharro, S. Panzeri, Synergy and redundancy in dual decompositions of mutual Information gain and information loss, Entropy 19 (71) (2017) 1–29. DOI:10.3390/e19020071.

[13] F. Kenney, E. S. Keeping, Mathematics of statistics, Pt.2, D. Van Nostrand Company, Inc., London, 1951.
